# Triacylglycerol stability limits futile cycles and inhibition of carbon capture in oil-accumulating leaves

**DOI:** 10.1101/2023.09.12.557462

**Authors:** Brandon S. Johnson, Doug K. Allen, Philip D. Bates

## Abstract

Engineering plant vegetative tissue to accumulate triacylglycerols (TAG, e.g., oil) can increase the amount of oil harvested per acre to levels that exceed current oilseed crops. Engineered *Nicotiana tabacum* (tobacco) lines that accumulate 15% to 30% oil of leaf dry weight resulted in starkly different metabolic phenotypes. In-depth analysis of the leaf lipid accumulation and ^14^CO_2_ metabolic tracing mechanistically described metabolic adaptations to the leaf oil engineering. An oil-for-membrane lipid tradeoff in the 15% oil line (referred to as HO) was surprisingly not further exacerbated when lipid production was enhanced to 30% (LEC2 line). The HO line exhibited a futile cycle that limited TAG yield through exchange with starch, altered carbon flux into various metabolite pools and end products, and suggested overlapping pathways of the glyoxylate cycle and photorespiration that limited CO_2_ assimilation by 50%. In contrast, inclusion of the LEC2 transcription factor in tobacco improved TAG stability, alleviated the TAG-to-starch futile cycle, and recovered CO_2_ assimilation and plant growth comparable to wild type but with much higher lipid levels in the leaves. Thus, the unstable production of storage reserves and futile cycling limit vegetative oil engineering approaches. The capacity to overcome futile cycles and maintain enhanced stable TAG levels in LEC2 demonstrated the importance of considering unanticipated metabolic adaptations while engineering vegetative oil crops.

## INTRODUCTION

As world population continues to grow, so do societal needs for energy feedstocks and chemicals obtained from finite supplies of petrochemicals (Singh et al., 2021). An extensive amount of biofuels research is focused on leafy biomass (which is less than 5% lipid (Li-Beisson et al., 2013)) and involves the breakdown and fermentation of recalcitrant carbohydrate polymers (Himmel et al., 2007; McCann and Carpita, 2015) that is an energetically intensive process. Oleochemicals from natural sources are an alternative that have comparable energy density to petrochemicals and a range of chemical structures for use by the chemical industry (Hill, 2007). Oleochemicals derived from plant oils (e.g., triacylglycerols, TAG) are the most energy-rich product of photosynthetic carbon assimilation with over twice the energy density of sugars and can be utilized for food, feed, biofuels, or as feedstocks to the chemical industry for various products such as plastics, polymers, resins, lubricants, etc.; however, production of oil per acre is not comparable to cellulosic levels in plants. Dramatic increases in plant oil production on the same or less arable land are needed to meet the growing global demands. Though most plants accumulate significant amounts of oil in the seeds or fruit, seed oil accumulation occurs during the last 1/3^rd^ of the plant lifecycle and does not capitalize on available vegetative biomass production. Engineering leafy biomass crops to accumulate energy-dense lipids at oilseed levels would dramatically enhance renewable energy feed stock production.

Vegetative oil accumulation has been achieved to varied extents (Xu and Shanklin, 2016; Vanhercke et al., 2019), including 4.3 & 8% TAG in sugarcane stems and leaves, respectively (Parajuli et al., 2020); 3.3% in potato tubers (Liu et al., 2017); 8.4% in *Sorghum bicolor* (Vanhercke et al., 2019); 8.7% TAG in leaves of duckweed (Liang et al., 2023); 2.5% TAG in perennial ryegrass (Beechey-Gradwell et al., 2020); and 9% in the leaves of the model plant Arabidopsis (Fan et al., 2014) as a percent of dry weight. None of these compare to that reported in tobacco leaves of ∼15-30% oil (Vanhercke et al., 2014; Vanhercke et al., 2017). Using a “Push-Pull-Protect” strategy resulted in ∼15% leaf dry weight as TAG (high oil, HO) lines (Vanhercke et al., 2014). The “Push” was generated by expression of the *Arabidopsis thaliana* WRINKLED1 (WRI1) transcription factor, which increased fatty acid biosynthesis, the *A. thaliana* DIACYLGLYCEROL ACYLTRANSFERASE 1 (DGAT1) provided the “Pull” of acyl groups into TAG, and the oil-body packing protein *Sesamum indicum* OLEOSIN provided a hypothetical “Protect” function. The HO lines accumulated 15% dry weight oil but were severely stunted in growth. When the *A. thaliana* LEAFY COTYLEDON 2 (LEC2) transcription factor (a regulator of embryogenesis) under control of the *A. thaliana* SAG12 senescence-inducible promoter was expressed in the HO background, leaf TAG content increased to ∼30% dry weight and surprisingly recovered plant growth near wild-type (WT) level (Vanhercke et al., 2017).

One of the perceived challenges in engineering storage oil production is to avoid compromising essential membrane lipid biosynthesis (Bates, 2016) because of significantly overlapping fatty acid biosynthetic and lipid assembly steps. However, the requirement for membranes in leaves presents an opportunity to co-opt some capacity with additional genes or transcription factors to enhance oil storage in vegetative tissues. With fatty acid biosynthesis in the plastid enhanced by AtWRI1 and TAG synthesis by AtDGAT1 in the endoplasmic reticulum of the engineered lines, the required enzymatic steps to produce TAG in leaves are in place (Li-Beisson et al., 2013). Thus, the successful accumulation of lipids in leaves depends on plant metabolism accommodating the engineered increased push and pull of carbon through endogenous metabolic networks into TAG. Understanding how tobacco leaf metabolism adapts to the engineered Push and Pull of carbon into leaf lipids in both the HO and LEC2 lines is a crucial part of the Design-Build-Test-Learn cycle to further enhance lipid metabolism in any species including more productive biomass crops (Pouvreau et al., 2018).

Isotopic labeling studies can track altered acyl flux resulting from lipid metabolic engineering and identify endogenous bottlenecks that affect the accumulation of the desired lipid products (Eccleston and Ohlrogge, 1998; Bates and Browse, 2011; Bates et al., 2014; Yang et al., 2017; Regmi et al., 2020). Previously [^14^C]acetate tracing of lipid metabolism in isolated leaf disks exposed to constant light indicated that acyl flux in HO was diverted away from photosynthetic membrane production to produce TAG with concomitant increased lipid turnover (Vanhercke et al., 2017; Zhou et al., 2020). It remained unclear if the enhanced lipid breakdown was partially induced by incubation of excised leaf disks in constant light, and further, what were the consequences of enhanced synthesis/turnover of lipids on carbon partitioning into other metabolic end products such as starch, protein, cell walls, or aqueous-soluble metabolic intermediates. The production and turnover of lipid and starch are crucial to optimal photosynthetic performance and plant growth (Huber and Hanson, 1992; Yu et al., 2018; Koper et al., 2021), and the presence of futile cycles can contribute to changes in growth phenotypes observed in tobacco (Edwards et al., 1999; Baud et al., 2009; Fan et al., 2019; Zhai et al., 2021). ^14^CO_2_ has been successfully utilized in tobacco to track carbon uptake and partitioning into different metabolic products (Olesinski et al., 1995; Häusler et al., 1998; Oparka et al., 1999). In this study, WT, HO, and LEC2 tobacco lines were ^14^CO_2_ pulse-chased, in planta, to examine the effects of leaf oil engineering on resource partitioning of photosynthetically fixed carbon into various metabolic products.

## RESULTS

### ^14^CO_2_ tracing indicates TAG in HO is metabolically dynamic rather than a stable end-product

To understand the effects of the diurnal cycle on lipid production and turnover, 62-day old WT and HO tobacco plants were pulsed with ^14^CO_2_ for 1.5 hours and allowed to grow for two day/night cycles. At the end of the second night, the plants were maintained in the dark for an additional 61 hours (Supplemental Fig. S1). The constant dark period was used to assess if leaf TAG turnover may be induced by energy starvation resulting from a lack of light which might occur during harvest and transport of green leaves for biofuel production. The ^14^C incorporation into total lipids, polar lipids (membrane lipids), and TAG was tracked across the 88 hr time course (Supplemental Fig. S1).

WT leaves initially produced more ^14^C labeled lipids than HO and reached a stable level over the time course; however, total levels of ^14^C lipids in HO were much more dynamic and decreased over time (Supplemental Fig. S1A). In WT, most of the ^14^C was associated with membrane lipids and ∼1% in TAG (Supplemental Fig. S1B, C), consistent with previous leaf labeling and may reflect TAG involvement in leaf lipid homeostasis (Xu and Shanklin, 2016; Karki et al., 2019; Zhou et al., 2020). The HO line accumulated less labeled polar lipids and more TAG than WT, suggesting carbon for membrane lipids is re-routed to TAG in HO (Supplemental Fig. S1B), consistent with [^14^C]acetate labeling of HO leaf disks (Zhou et al., 2020) and indicating good agreement between excised leaf disk and whole plant metabolic labeling experiments. TAG in HO, reached 17% of labeled lipids 3 hrs after the pulse before declining to 3-4% at the end of the chase (Supplemental Fig. S1C), indicating that HO leaves remobilize carbon from TAG. This initial experiment demonstrated the feasibility of utilizing ^14^CO_2_ to trace differences in carbon metabolism between WT and oil-producing tobacco plants over extended periods; therefore, the approach was extended to analyze more metabolites and include the LEC2 line.

### Leaves in mid-development show the greatest differences in oil accumulation

A development study was performed to determine the best stage in plant development for ^14^CO_2_ metabolic labeling based on: (1) maximal differences in lipid accumulation between WT, HO, and LEC2 lines; and (2) plant size that (a) maximized the available leaf tissue area for sampling and (b) that accommodated the most plants within a 65L chamber for ^14^CO_2_ labeling. Total lipid fatty acid content in leaf disks were directly converted to fatty acid methyl esters (FAMEs) and quantified by gas chromatography at 30-, 44-, and 64-days after sowing (Fig. 1A). The HO plants were smaller than WT while LEC2 plants more closely approached the size of WT (Vanhercke et al., 2017). In the relatively small 30-day plants, the HO leaves accumulated an average of approximately 600 µg FAMEs per leaf disk, nearly 3-fold more lipid content than WT, but not significantly different than LEC2 at this stage. The medium-sized 44-day old plants had the most significant difference in lipid accumulation between the lines, where LEC2 averaged approximately 900 µg (and up to 1200 µg) FAMEs per leaf disk. While the large 64-day old plants (similar age to plants in Supplemental Fig. S1) indicated the most oil in the LEC2 lines, they were too large to be used in replicate with the other lines in the 65L chamber. To confirm that changes in total lipid content of mid-size plants reflected leaf TAG production; we quantified the major neutral and polar lipids (Fig. 1B) and associated fatty acid composition (Supplemental Figure S2-S3) in each line from 40-day old plants. The differences between WT and HO polar lipids and TAG content indicated a large increase in HO TAG content (∼12060%) with a concomitant ∼43% decrease in galactolipids relative to WT. The further increase in LEC2 TAG over HO at 40 days after sowing was modestly compensated by an insignificant reduction in mean galactolipid and phosphatidylglycerol content (Fig. 1B). Thus, the LEC2 line greatly increased leaf oil content while maintaining comparable chloroplast membrane content. Based on plant size, leaf area, and differences in lipid accumulation, 40-45 days old plants were chosen for additional ^14^CO_2_ pulse-chase analysis.

**Figure 1.**
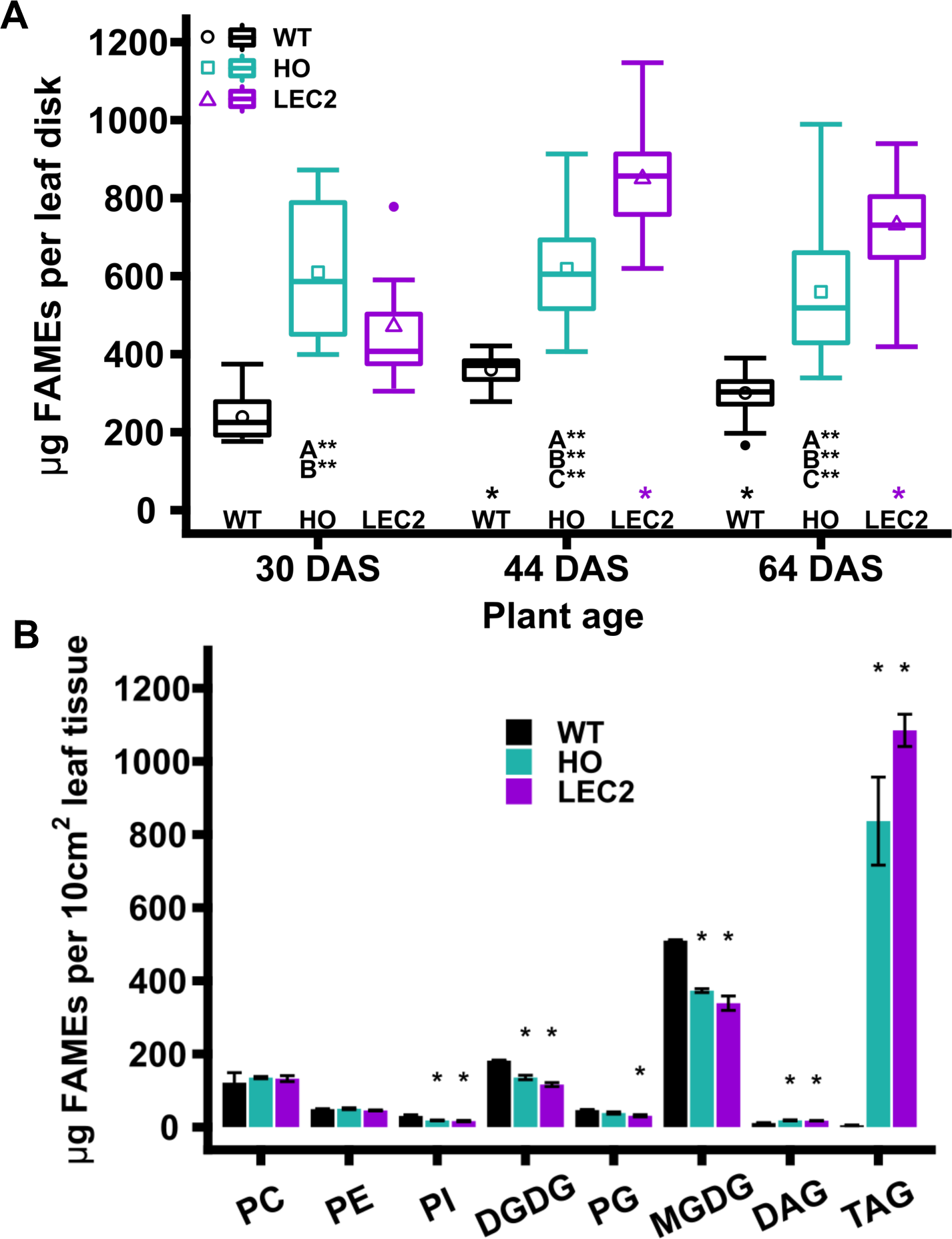
Lipid accumulation in WT, HO, and LEC2 tobacco leaves. A, Total leaf lipid accumulation at 30, 44, and 64 days after sowing (DAS). The box represents the interquartile ranges of 25-75%, the upper whisker represents the maximum value that is within 1.5 times over the 75th percentile and the lower whisker represents the minimum value that is within 1.5 times under the 25th percentile, the cross bar in the box represents the median, the shape represents the mean, and outliers are points outside of the range of the whiskers ( > 1.5 times interquartile range). Leaf disks collected (n) during study (30 DAS: n = 9-18; 44 DAS: n = 31-54; 64 DAS: n = 67-107). Significant differences between lines at each age are indicated by comparison, A: WT – HO; B: WT – LEC2; C: HO – LEC2 and significance differences within a line are indicated across the x-axis (Reference group = 30 days) P-values (< 0.05: *, < 0.01: **) were calculated by ANOVA and Tukey Honest Significant Differences test for multiple comparisons. All leaf tissue normalized to 254.5 mm2. B, Glycerolipid analysis for wild-type (WT) and oil-accumulating (HO, LEC2) tobacco. A cork bore (18 mm diameter) was used to collect leaf disks from 40-day old plants. Five leaf disks were collected from various leaves at random from a single plant and combined for a single sample. Bars represent the mean (n = 3, except LEC2 DAG which has n = 2) and error bars represent mean +/− one standard error. Asterisks above HO and LEC2 bars indicate significant differences (p-value < 0.05) compared to WT (reference group). No significant differences between HO and LEC2.

### Differential lipid accumulation during a 145 hr ^14^CO_2_ pulse-chase labeling of WT, HO, and LEC2 tobacco

A 2 hr ^14^CO_2_ pulse and chase of 145 hrs was utilized to analyze the partitioning of photosynthetically fixed carbon into major metabolites between the tobacco lines. Fig. 2A-B demonstrates the labeling chamber and the sampling protocol implemented to minimize differences in individual leaf development between replicate plants and maximize the total leaf area available for sampling. Eight time points were collected across three sequential horizontal leaves on each replicate plant. Two wild-type replicate plants and three replicate plants of both HO and LEC2 at 45 days old were able to fit in the 65L labeling chamber together. At the end of the pulse (designated time 0 hr), leaf disks were collected from three sequential leaves per plant (n = 12-18 at 0 hr), and from one leaf per plant at later time points (n = 4-6) (Fig. 2B). Leaf disk biomass was separated into five major fractions: aqueous soluble metabolites, total protein, cell wall, starch, and total lipids (Fig. 2C). In addition, the lipid fraction was further divided between TAG and polar lipids, and the aqueous fraction was further split out into sugars, organic acids, and amino acids. Plants appeared healthy during the entire experiment and did not show signs of growth defects from sampling (Fig. 2D). The lines varied in lipid accumulation over the 145-hour chase time course (Fig. 2E). WT increased lipid mass from 301 µg to 486 µg FAME per leaf disk over development. At the end of the pulse, leaf disks collected from HO and LEC2 had a similar amount of lipids which was ∼1.3-1.4 fold greater than WT. Though HO started with more total lipid mass than WT, total FAME per leaf disk in HO only increased slightly from 446 µg to 500 µg FAME over the 145 hr time course. By 100 hrs after the pulse, total WT lipid mass reached the level of HO and both did not change further over the next 45 hrs. During the time course, LEC2 lipid mass per leaf disk increased significantly from 406 µg FAME at hour 0 to 1133 µg FAME at hour 145, an increase of nearly 300% in seven days. Thus, the plants that were ^14^CO_2_ labeled grew well and had changes in carbon metabolism leading to differential lipid accumulation over the time course.

**Figure 2.**
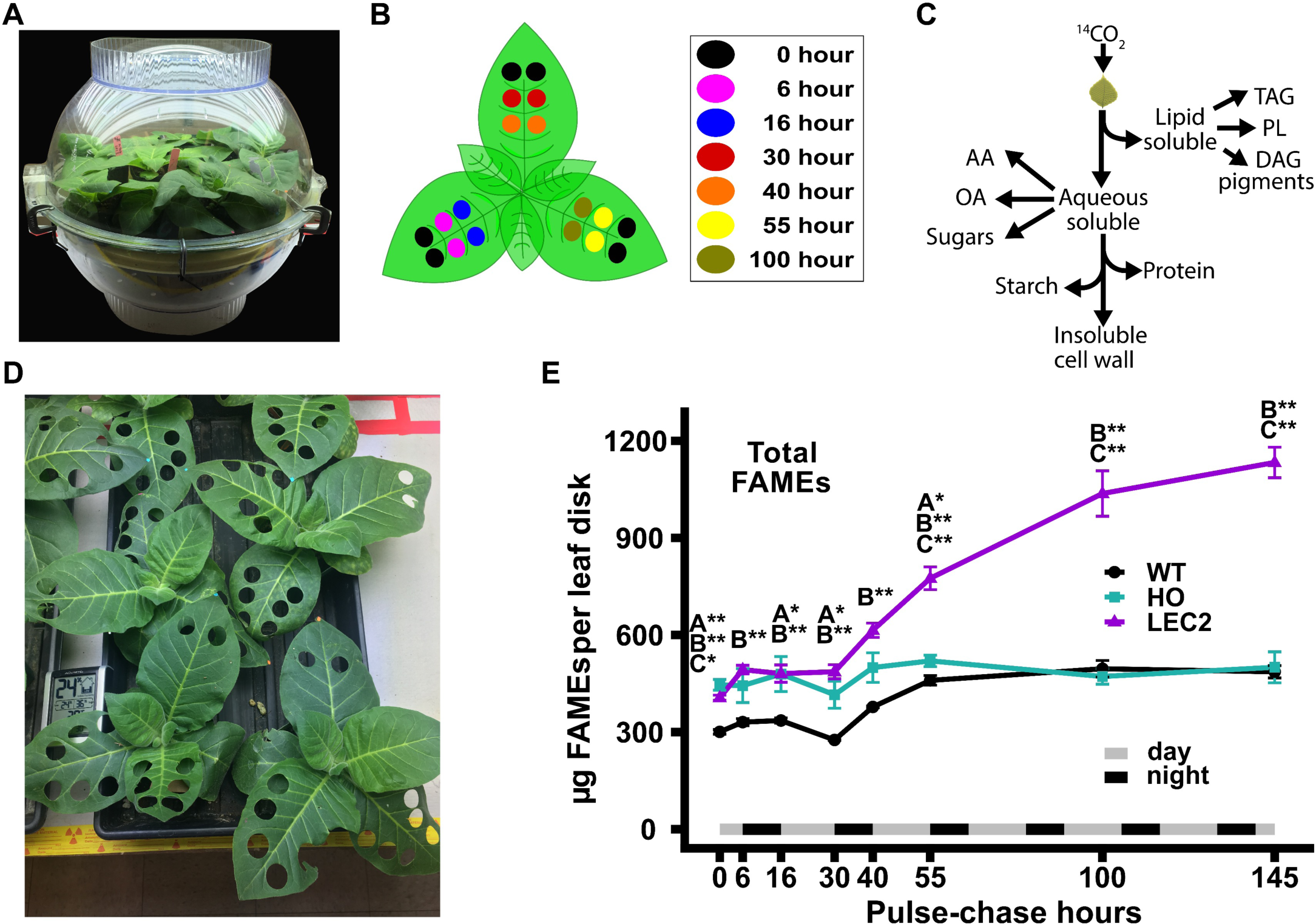
^14^CO_2_ pulse-chase experimental design. (A) Labeling chamber, ∼ 65 L volume, with two small fans inside to increase circulation of 1 mCi ^14^CO_2_ during 2-hour pulse. (B) ^14^CO_2_ pulse-chase sampling protocol: Two WT and three HO and LEC2 plants were sampled at eight time points distributed across three sequential horizontal leaves. Leaf disks were collected from all the leaves at the end of the pulse: 0 hr, *n*_(WT)_ = 12, *n*_(HO,LEC2)_ = 18 disks. At hrs 6-100, duplicate leaf disks were collected to represent each leaf number at single time, in an alternating pattern as indicated by colors; *n*_(WT)_ = 4, *n*_(HO,LEC2)_ = 6. Hour 145 is not shown in (B) since leaf disks were collected randomly where space allowed. Leaf disks were collected by a 14 mm diameter cork bore, ∼ 154 mm^2^ leaf tissue. (C) Metabolite extraction by sequence of fractionations. (D) Plants growth during time course. Plants continued growing and appeared healthy after leaf disk collection with the picture taken after the fourth time point (30 hours after pulse). (E) Total lipid mass in leaf disks from the 145 hr ^14^CO_2_ pulse-chase. The unlabeled lipid mass measured as fatty acid methyl esters (FAME) from lipid extracts in Figure 5F. All data points are mean ± SE. Asterisks indicating significant differences between lines, A: WT – HO; B: WT – LEC2; C: HO – LEC2 (ANOVA and Tukey Honest Significant Differences test for multiple comparisons) p-value 0.05 – 0.01 = *, < 0.01 = **.

### Metabolic partitioning of ^14^CO_2_ into major metabolite fractions differed significantly between lines

At the end of the pulse (Fig. 3A, 0 hr), total ^14^C accumulation indicated significant differences in ^14^CO_2_ assimilation between each tobacco line. Total ^14^C, measured as disintegrations per minute (DPM) per leaf area, in WT was over two-fold greater than HO, and more than 25% greater than LEC2. Total radioactivity decreased over the first 55 hours of the chase period before mostly leveling off, indicating partitioning into storage reserves that were less significantly turned over. By 145 hrs, the total ^14^C in each line was less than half the value at the end of the pulse (Fig. 3A). The initial decrease in total ^14^C per leaf area most likely represented the export of photosynthetically fixed carbon from the leaf to other tissues. CO_2_ converted into different metabolite fractions was quantified as a percentage of the total radioactivity across the time course (Fig. 3B-F). The relative proportion of carbon partitioning to aqueous soluble metabolites was not initially significantly different at chase hour 0 between any of the tobacco lines (Fig. 3B), however, the oil accumulating lines had a more rapid efflux of ^14^C out of the aqueous fraction as the aqueous soluble metabolites were converted to other metabolites during the chase period. By hour 55, WT, HO, and LEC2 were similar in the percentage of ^14^C in the aqueous fraction and protein (Fig. 3C). At the beginning of the chase, ^14^C partitioning to the cell wall fraction in HO (21.7%; Fig. 3D) was approximately 1.5-fold greater than WT (14.1%) and more than 1.9-fold greater than LEC2 (11.2%). HO had a higher percent of ^14^C associated with the cell wall fraction than both WT and LEC2 throughout the experiment, though all lines exhibited similar ^14^C increases in cell wall over the time course (Fig. 3D).

**Figure 3.**
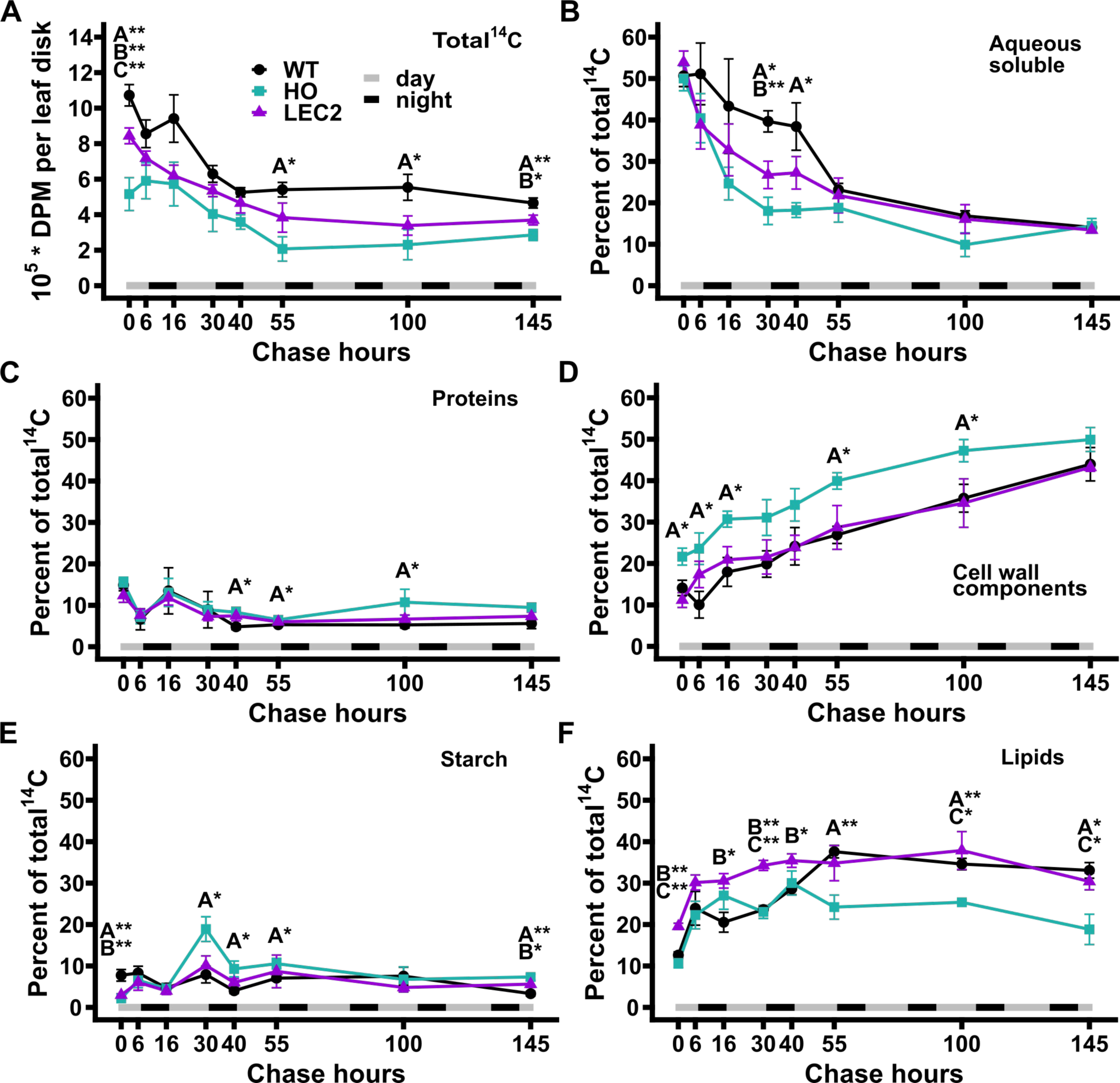
Total radioactivity and label distribution among major metabolite fractions from 145 hr ^14^CO_2_ pulse-chase of 45-day old WT, HO, and LEC2 lines. 2 hr pulse with 1 mCi ^14^CO_2_, and chase of 145 hrs. 0 hr is the end of the pulse. (A) Total ^14^C accumulation in ∼ 154 mm^2^ leaf disks. (B-F) Partitioning of ^14^C to metabolites in the aqueous, protein, cell wall, starch, and lipid phases expressed as a percent of total ^14^C. Leaf disk collection just prior to lights off and lights on for hours 6, 16, 30, 40, and 55, then every other day for hours 100 and 145. At hour 0, *n*_(WT)_ = 12, *n*_(HO,LEC2)_ = 18 disks. At hours 6-145, *n*_(WT)_ = 4, *n*_(HO,LEC2)_ = 6 disks. All data points are mean ± SE. Asterisks indicating significant differences between lines, A: WT – HO; B: WT – LEC2; C: HO – LEC2 (ANOVA and Tukey Honest Significant Differences test for multiple comparisons) p-value 0.05 – 0.01 = *, < 0.01 = **.

At the end of the pulse (time 0 hr), WT contained significantly more ^14^C within starch (2.6-3.6-fold more) than the high oil lines (Fig. 3E, 0 hr). Within the first 6 hours of the chase, labeled starch increased more rapidly within the oil lines. Then labeled starch decreased in all lines between 6 and 16 hrs, consistent with starch turnover diurnally. Over the second light period (chase hours 16-30) the percent of total ^14^C in starch increased in all lines; though HO was significantly greater, reaching 18.9% of total ^14^C, compared to only 7.9% and 10.1% for WT and LEC2, respectively. At the end of the second night (chase hr 40) diurnal starch turnover had resulted in a decrease of ^14^C starch by approximately 50%. A dampened increase in labeled starch was also apparent during the 3^rd^ day period (chase hours 40-55) and decreased between 55 and 145 hours of the chase in all lines. By the end of the chase, HO contained more labeled starch (7.3%) than LEC2 (5.6%) or wild type (3.3%), distinct from the end of the pulse (0 hour) where WT was 2.6-3.6-fold higher than the oil lines.

The partitioning of fixed ^14^C into total lipids revealed unique differences between each line (Fig. 3F). At hour 0, LEC2 allocated 19.6% of total ^14^C to lipids, whereas WT and HO were 12.6% and 10.7%, respectively. LEC2 maintained a higher percentage of labeled lipids over both WT and HO until hour 55, when total labeled lipid content on a percent basis was similar in LEC2 and WT and remained so for the duration of the time course. During the first 55 hours of the time course, ^14^C lipid labeling within HO was changing more extensively to offset starch levels and potentially accommodate diurnal carbon needs. By the end of the time course, the fraction of ^14^C accumulated in HO total lipids was significantly less than WT and LEC2. In summary, total ^14^CO_2_ uptake was decreased in the oil lines compared to WT, with HO having the largest disparity of less than ½ the fractional level of lipid relative to WT. LEC2 carbon uptake was measured to be 78.7% of WT, indicating a significant improvement in carbon assimilation relative to HO. Both HO and LEC2 exhibited significant differences in partitioning of fixed ^14^C from WT and each other, particularly in the cell wall, lipid, and starch fractions indicating metabolic alterations in addition to lipid accumulation in HO and LEC2 lines.

### Aqueous metabolite fractions show the greatest differences in organic acid labeling

To further investigate the changes in carbon partitioning between WT and the oil-producing lines, the aqueous metabolite fraction (Fig. 3B) was split into neutral, anionic, and cationic aqueous metabolite fractions essentially composed of soluble sugars, organic acids, and amino acids, respectively (Fig. 4). At the end of the ^14^CO_2_ pulse (0 hr) the organic acid portion (Fig. 4A) contained more ^14^C than other aqueous fractions in all lines with WT accumulating the most (30%, 27%, and 18% in WT, LEC2 and HO, respectively) as a percent of total radioactivity (Fig. 4B). The amount of labeled organic acids decreased more rapidly in both oil-producing lines between 0-40 chase hours than WT (Fig. 4B), suggesting more rapid turnover of organic acid pools in oil lines than WT, less synthesis of labeled organic acids during the chase period from turnover of other labeled pools in the oil lines and possibly larger organic acid concentrations in the vacuoles within WT that were labeled. By 55 hrs into the chase period, the labeled organic acid levels in the WT were similar to the oil lines (Fig. 4B). The percent of total radioactivity in the soluble sugar fraction was similar among the lines and decreased from ∼13-14% at the end of the pulse (Fig. 4C). The amino acid portion was the least labeled in all lines (Fig. 4E), with total radioactivity in the HO line highest on average and significantly higher than WT at 6 of the 8 time points (Fig. 4F). The free amino acid fraction indicated larger differences between HO and the other two lines (Fig. 4F) than the total protein fraction (Fig. 3C), suggesting that the chase turnover of labeled metabolites led to a higher flux through free amino acids pools that was not directly involved in protein synthesis.

**Figure 4.**
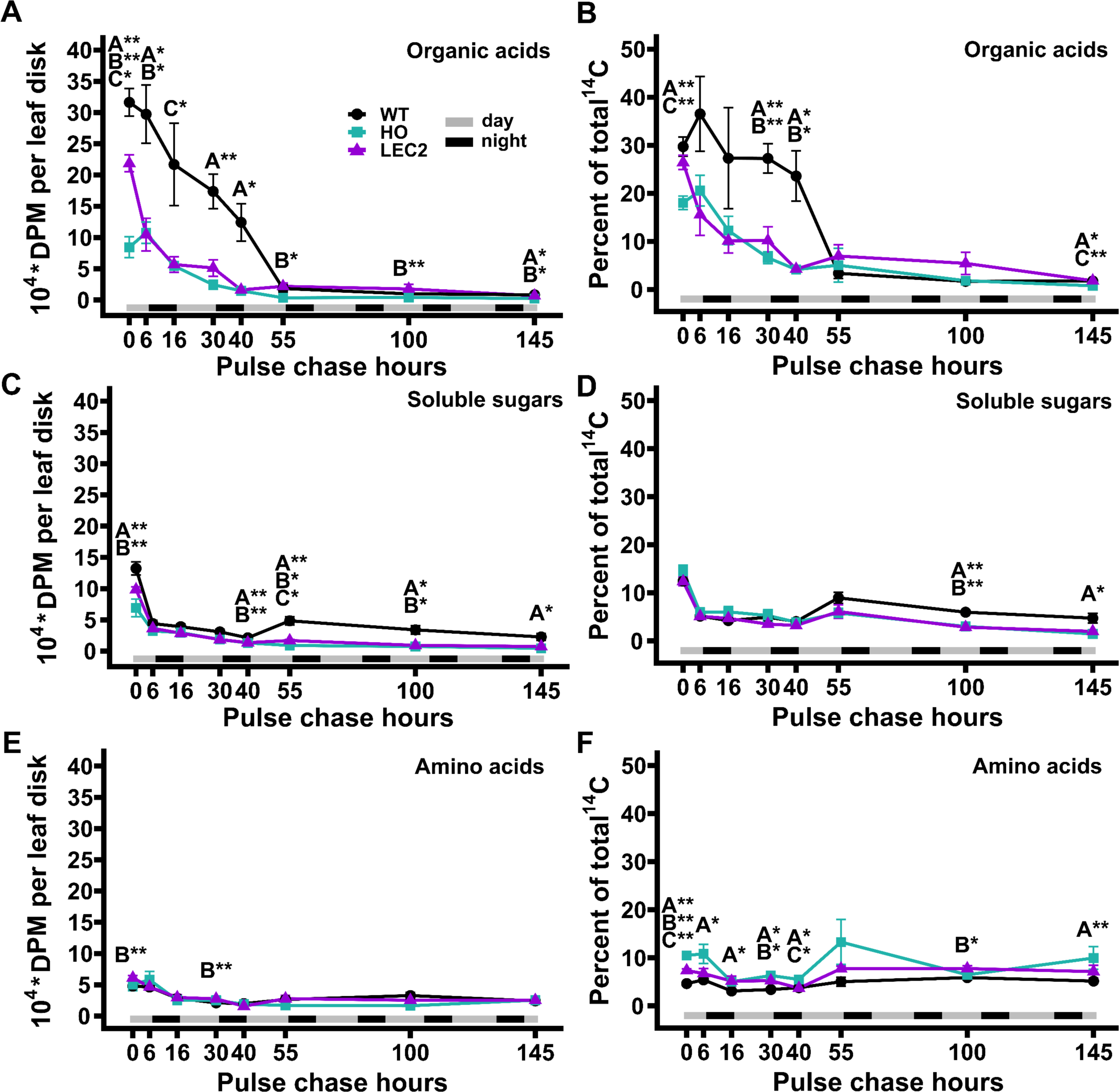
Quantitation of radiolabeled sugar, organic acid, and amino acid fractions from the 145 hr ^14^CO_2_ pulse-chase of WT, HO and LEC2 tobacco lines. A) Sugar, B) organic acid, C) amino acid fractions expressed as a percent of sum of [^14^C]aqueous fraction from Figure 5B. All data points are mean ± SE. Asterisks indicating significant differences between lines, A: WT – HO; B: WT – LEC2; C: HO – LEC2 (ANOVA and Tukey Honest Significant Differences test for multiple comparisons) p-value 0.05 – 0.01 = *, < 0.01 = **.

### Distinct patterns of labeled lipid fluxes in each tobacco line

The lipid fraction (Fig. 3F) was separated into neutral lipids and polar lipids by thin-layer chromatography and radioactivity in each lipid class was quantified by phosphor imaging (Supplemental Fig. S4). Similar to the initial ^14^CO_2_ labeling experiment (Supplemental Fig. S1) the WT radiolabeled lipid fraction was predominantly composed of polar lipids (e.g., membrane lipids) (Fig. 5A, C, E) with lesser contribution from diacylglycerol, free sterols, steryl esters, and pigments (Supplemental Fig. S5). TAG represented less than 1% of the labeled lipids in WT (Fig. 5B, D, F). The total radioactivity per leaf disk in WT polar lipids reached a maximum at 6 hours of the chase and remained roughly constant over the time course (Fig. 5A); though total fixed ^14^C was decreasing during the same initial period of the chase (Fig. 3A), thus resulting in increased polar lipid percentage as a fraction of total radioactivity of 24.2% (Fig. 5C). When considering the partitioning of ^14^C within lipids, the label in WT polar lipids was unchanged indicating polar lipid composition was maintained (Fig. 5E). The results indicated that WT membrane lipids synthesized during and shortly after the pulse were stable over the time course.

**Figure 5.**
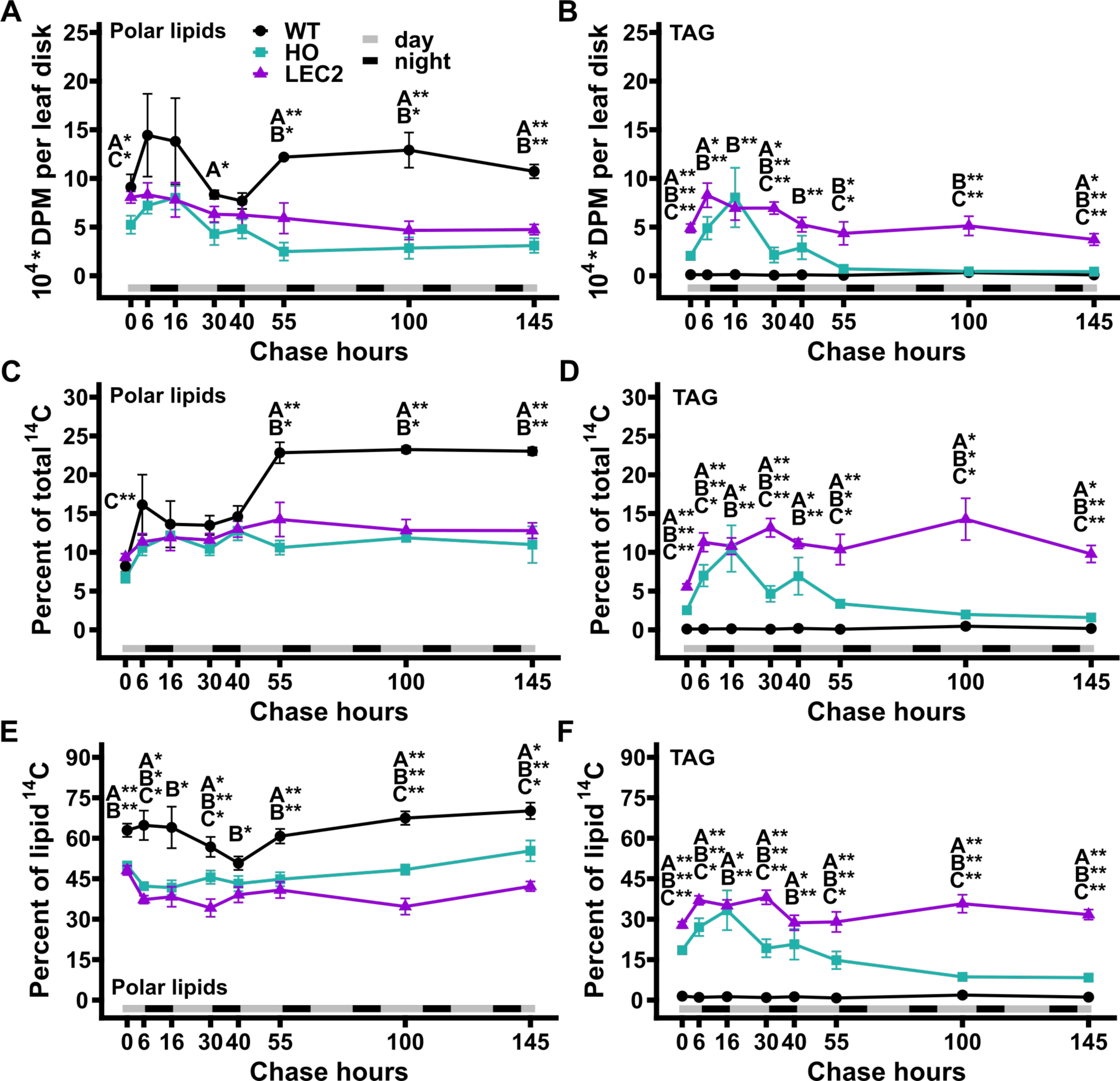
Fractionation and quantitation of radiolabeled lipids from the 145 hr ^14^CO_2_ pulse-chase of WT, HO and LEC2 tobacco lines. The total lipid extract in Figure 5F fractionated by TLC into: A;C;E) Polar lipids; B;D;F) TAG. All data points are mean ± SE. Asterisks indicating significant differences between lines, A: WT – HO; B: WT – LEC2; C: HO – LEC2 (ANOVA and Tukey Honest Significant Differences test for multiple comparisons) p-value 0.05 – 0.01 = *, < 0.01 = **.

As expected from the lipid mass quantification (Fig. 1A, 4E) the high oil lines both accumulated significantly more ^14^C-TAG (Fig. 5B, D, F) and less labeled polar lipids (Fig. 5A, C, E) than WT by the end of the pulse. However, each oil line exhibited distinct patterns of ^14^C lipid flux. In HO, total radioactivity per leaf disk and percent of total ^14^C for both polar lipids and TAG increased from the end of the pulse through the remaining day and first night (0-16 chase hrs, Fig. 5A-D). After 16 hrs of chase the total ^14^C per leaf disk for both polar lipids and TAG (Fig. 5A-B) rapidly decreased during the second day (16-30 hrs), increased slightly during the second night (30-40 chase hrs), then rapidly decreased again during the third day (40-55 chase hrs). At 55 hrs of chase the total ^14^C per leaf disk in HO polar lipids was about half of the maximum at 16 hrs, however total ^14^C in TAG had decreased 94% from the 16 hr maximum (Fig. 5A-B). From 55-145 hrs the remaining total radioactivity in HO polar lipids was stable while TAG continued to decrease (Fig. 5A-B). As a percent of total fixed ^14^C (Fig. 3A), the total polar lipid labeling fluctuations were dampened and became near constant from 16-145 hrs (Fig. 5C). However, HO TAG as a percentage of total ^14^C was similar to the total TAG ^14^C per leaf disk demonstrating a dynamic pattern (Fig. 5D). The results indicated that labeled TAG in HO was not stable and ^14^C flux through the TAG pool was dependent on diurnal cycles, with ^14^C in TAG decreasing during the day and increasing at night. In addition, ^14^C flux through the HO polar lipid pools only partially mirrored the TAG pool. In LEC2, initial labeling of polar lipids at the end of the pulse was 35% and 43% more than HO as total ^14^C per leaf disk and percent of total ^14^C, respectively (Fig. 5A, C), while LEC2 initial TAG labeling was more than double that of HO by both measures (Fig. 5B, D). Total ^14^C per leaf disk for both polar lipids and TAG of LEC2 reached a maximum by the end of the first day (6 chase hrs), and slowly decreased over the time course by 43% and 50%, respectively (Fig. 5A-B). When considered as a percent of total fixed ^14^C, both polar lipids and TAG in LEC2 were roughly constant over the time course (Fig. 5C-D). Together these results indicated that both TAG and membrane lipids were more stable in LEC2 than HO.

Membrane lipids and TAG are produced through a complex metabolic network in which intermediates (e.g., fatty acids, diacylglycerol (DAG)) are exchanged between membrane lipids and TAG (Bates, 2016, 2022). Therefore, changes in relative lipid class radioactivity were examined over the pulse-chase period (Fig. 5E-F). In WT most lipid radioactivity was partitioned into polar lipids with very little in TAG which remained near constant over the time course (Fig. 5E). In both the oil producing lines the percent of ^14^C initially incorporated into polar lipids decreased over the first 6 hours of chase (Fig. 5E) and was accompanied by an increase in TAG (Fig. 5F) consistent with fatty acids (and possibly DAG) transiently incorporated into PC prior to accumulation in TAG (Bates, 2016; Zhou et al., 2020). In HO the percent of total lipid ^14^C-TAG decreased from 16-145 chase hours, and polar lipids increased slightly (Fig 5. E-F) suggesting the turnover of HO TAG (Fig. 5B, D) that consequently increases percent polar lipid levels (Fig. 5A, C) but may also indicate some transfer of ^14^C from TAG to polar lipids during the chase. In LEC2, the relative labeling of polar lipids and TAG from 6-145 hrs was roughly constant further supporting the stability of TAG in the LEC2 line (Fig. 5E-F).

### LEC2 alleviates a TAG-starch futile cycle present in HO leading to higher TAG content

The relative accumulation of TAG and starch as both total radioactivity (DPM) and percent of total ^14^C in WT, HO, and LEC2 across the 145 hr time course was compared (Figure 6). At 45-days, diurnal cycling of ^14^C-starch was apparent in all lines (Fig. 6A-F); however, the HO line exhibited an inverse pattern of TAG diurnal cycling (Fig. 6C-D). During the first night (6-16 hrs) labeled starch decreased and labeled TAG increased suggesting the breakdown of starch at night fuels HO TAG biosynthesis. Surprisingly ^14^C-TAG decreased concomitant with an increase in ^14^C-starch during the second day (16-30 hrs) indicating the turnover of carbon from TAG to indirectly fuel starch production in HO, which was not apparent in other lines. The pattern repeated though with more dampened response as ^14^C intermediates of starch and TAG metabolism were siphoned off for other products during the extended chase. While some diurnal cycling of ^14^C-starch was apparent in LEC2 (Fig. 6E-F), the amplitude at 16-40 hrs was less than HO, and there was little change in ^14^C-TAG content. The larger changes in both labeled starch and TAG in HO suggest a TAG-starch futile cycle which was alleviated in the LEC2 line. We hypothesize that examples of turnover may be commonplace in engineered systems that fail to meet expectations; however evidence for futile cycling requires dynamic labeling or other specific methods that can be difficult to obtain in practice.

**Figure 6.**
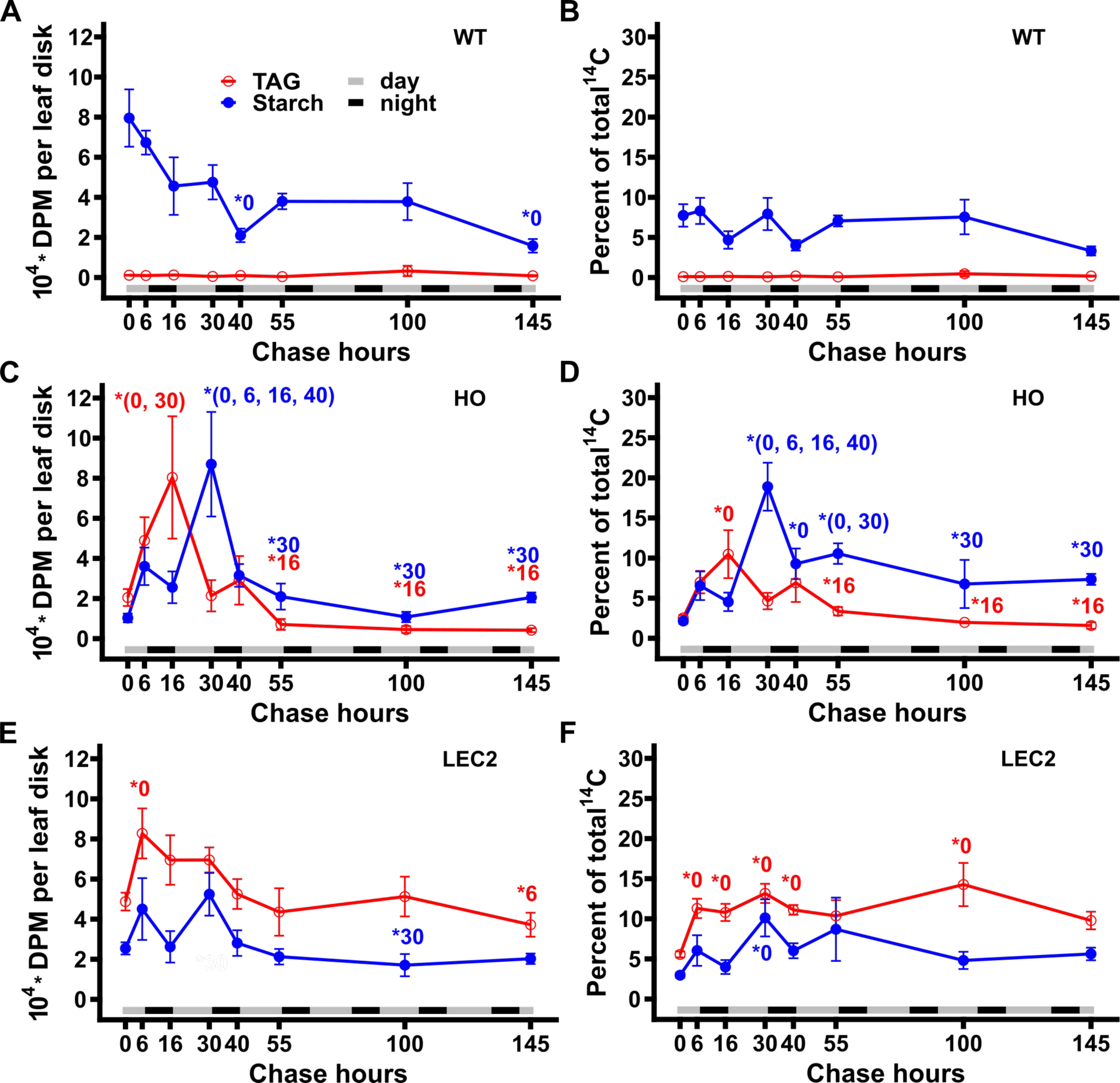
Comparison of radiolabeled TAG and starch from the 145 hr ^14^CO_2_ pulse-chase of WT, HO and LEC2 tobacco lines. Data obtained from Figures 4E and 7B;D and is represented as both total radioactivity in starch (blue) and TAG (red) (A, C, E), and as starch and TAG as a percent of total radioactivity (B, D, F). A-B, WT. C-D, HO. E-F, LEC2. All data points are mean ± SE. Asterisks indicating significant differences between lines, A: WT – HO; B: WT – LEC2; C: HO – LEC2 (ANOVA and Tukey Honest Significant Differences test for multiple comparisons) p-value 0.05 – 0.01 = *, < 0.01 = **.

## DISCUSSION

In this study, we investigated carbon partitioning in engineered tobacco with ∼15% oil in leaves (HO line), and a second line (LEC2 line) that additionally co-expresses *AtLEC2* in the HO background resulting in increased plant growth and accumulating significantly more oil (∼30% of dry weight) in leaves (Vanhercke et al., 2014; Vanhercke et al., 2017). Through quantitative assessment of the lipid phenotype and isotopic tracing into biomass components and pathway intermediates we demonstrate: 1) TAG within HO leaves can be remobilized out of storage pools during an energy deficit of extended darkness (Supplemental Fig. S1); 2) a period of rapid accumulation of leaf oil in LEC2 leaves from (30-44 DAS) that was absent from HO and is consistent with the delayed expression of the senescence-inducible promoter for *AtLEC2* (Fig. 1A); 3) oil accumulation in HO that coincides with reduced photosynthetic membrane lipid production, but increased TAG content of LEC2 that does not result in additional reductions in photosynthetic membrane lipids (Fig. 1B, Supplemental Fig. 2-3); and 4) differences in carbon assimilation and partitioning to lipids, aqueous-soluble, protein, starch, and cell wall components (Figs. 2-6) through a quantitative ^14^CO_2_ pulse-chase labeling design over a ∼7-day period (145 hours).

### ^14^CO_2_ labeling indicates differences in carbon assimilation and partitioning between tobacco lines

The ^14^CO_2_ pulse-chase labeling study revealed that leaves with enhanced TAG accumulation had significant alterations in central carbon metabolism, most prominently in the HO line. The HO line fixed approximately half as much ^14^CO_2_ as WT and exhibited significantly less ^14^C partitioned to starch (Fig. 3E, hr 0), more to cell wall (Fig. 3D, hr 0), and increased labeling in the amino acid fraction with concomitant decrease of ^14^C in the organic acid fraction compared to both WT and LEC2 (Fig. 4, hr 0). ^14^C in lipids initially did not differ significantly between HO and WT though LEC2 was nearly double the others (Fig. 3F, hr 0); 5). Over the chase period each line had a unique pattern of carbon partitioning as the metabolites labeled during the pulse were converted to downstream products (Figs. 3-6). Thus, the effect of the first iteration of leaf oil engineering (i.e., HO line) impacted more than fatty acid synthesis and TAG accumulation, it affected total CO_2_ fixation and carbon partitioning with consequences on other aspects of metabolism and biomass production. The inclusion of LEC2 did not further magnify the differential partitioning of HO, but instead partially compensated for the lost carbon assimilation and exhibited metabolism more consistent with WT. The results indicate that changes in growth and metabolic flux to accumulate oil have a widespread impact on central metabolism that exceeds fatty acid synthesis and lipid assembly and may support several intriguing hypotheses.

### Hypothesis 1: FA β-oxidation and glyoxylate cycle coinciding with photorespiration in leaf peroxisomes may reduce photorespiratory flux and CO_2_ assimilation in HO

The flux of ^14^C through a TAG-starch futile cycle in HO leaves (Fig. 6C-D) would involve the mobilization of carbon from TAG through β-oxidation and result in acetyl-CoA that could be used in the glyoxylate cycle or as a precursor of other storage reserves (such as starch through the glyoxylate cycle and gluconeogenesis). In a leaf, the production of glyoxylate occurs significantly through photorespiration (Fig. 7A) (Pan et al., 2020). The additional potential source of glyoxylate may contribute to the observed phenotype. ^14^CO_2_ enters leaf metabolism through photosynthesis when ribulose 1,5, bisphosphate carboxylase/oxygenase (RuBisCO) carboxylates ribulose 1,5 bisphosphate (RBP) that can be metabolically partitioned for fatty acids (Fig. 7A2) and TAG (Fig. 7A3). Photorespiration occurs when RuBisCO oxygenates RBP (Fig. 7A1). The pathways of photorespiration (Fig 7A, purple arrows), β-oxidation (Fig. 7A4), and glyoxylate cycle (Fig. 7A5, 7-8) have been extensively characterized and reviewed for many plant species including tobacco (Igamberdiev et al., 1995; Escher and Widmer, 1997; Igamberdiev and Kleczkowski, 2000; Graham, 2008; Bauwe et al., 2010; Tjellström et al., 2015; D’Andrea, 2016; Kelly and Feussner, 2016; Timm, 2020; Timm and Hagemann, 2020; Shi and Bloom, 2021).

**Figure 7:**
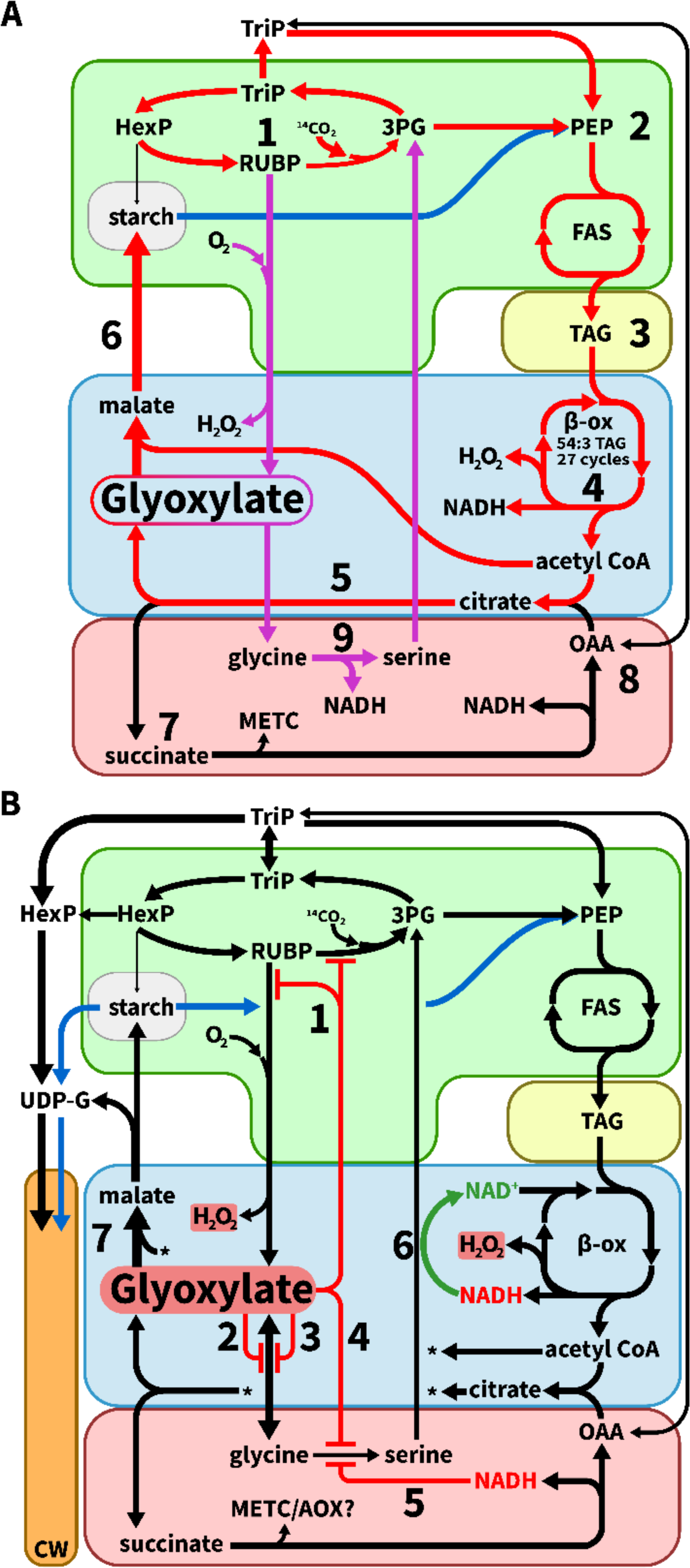
Reorganizing central metabolism to accommodate TAG turnover in tobacco leaves. A. TAG-starch futile cycle. Carbon flux through the TAG-starch futile cycle (red and blue arrows) and photorespiration (purple arrows) begins in the chloroplast (green) with (1) photosynthetic carboxylation (^14^CO_2_) or oxygenation (O_2_) of RUBP, respectively. (2) Calvin cycle partitioning of triose phosphates (triP) to fatty acid synthesis (FAS) is typically cytosolic, though expression analysis suggested a complete plastidial glycolytic pathway may be active to form PEP (Vanhercke et al., 2017). New photosynthetic carbon is partitioned to FAS for (3) TAG assembly while (4) β-oxidation degrades TAG that, releases H_2_O_2_, NADH, and old carbon as acetyl-CoA that combines with oxaloacetate (OAA) to form citrate in the glyoxylate cycle within the peroxisome (blue). The glyoxylate cycle converts (5) citrate to glyoxylate and succinate, with the former accepting an acetyl-CoA to form malate and the latter metabolized in the mitochondria (red) (cytosolic *aconitase* reaction step not shown). (6) Malate enters gluconeogenesis in the cytosol and some unknown intermediate is imported to the chloroplast to synthesize starch. (7) Succinate metabolism at complex II increases the redox potential in the mitochondrial electron transport chain (METC) during illumination in conjunction with the photosynthetic production of NADPH, while (8) replenishing oxaloacetate as the precursor for the glyoxylate cycle. Photorespiratory flux produces glyoxylate and H_2_O_2_ in the peroxisome and (9) NADH in the mitochondria via glycine decarboxylase complex (GDC). B. Inhibition via glyoxylate and NADH of photorespiratory flux stimulates cell wall synthesis. Glyoxylate is a potent inhibitor of (1)RuBisCO and RuBisCO activase, (2)glutamate:glyoxylate aminotransferase, (3)serine:glyoxylate aminotransferase, and (4) GDC, with GDC also inhibited by NADH (5). Photorespiratory flux to the chloroplast may be enhanced with the reducing potential generated within β-oxidation acting to reduce hydroxypyruvate to glycerate (6). Increased carbon partitioning to the cell wall (orange) in HO (Fig. 3D) may result from the inhibitory effects excess glyoxylate and NADH which redirects carbon flux from photorespiration to gluconeogenesis.

Efficient photorespiratory flux is necessary to maintain carbon assimilation (i.e., autotrophic metabolism) in atmospheric conditions and effectors such as NADH and glyoxylate, both produced within β-oxidation and glyoxylate cycle, respectively, can inhibit photorespiratory flux (Peterson, 1982; Havir, 1986; Escher and Widmer, 1997; Pritchard et al., 2002; Timm et al., 2012; Lu et al., 2014; Timm et al., 2016; Timm, 2020; Timm and Hagemann, 2020). High flux through FA β-oxidation and glyoxylate cycle may impair photosynthetic carbon fixation in several ways. Increasing cellular glyoxylate concentration can inhibit enzymes involved in photosynthesis (RubisCO and RuBisCO activase (Fig. 7B1)) and photorespiration (glutamate:oxoglutarate aminotransferase (GOGAT, Fig. 7B2), serine:glyoxylate aminotransferase (SGAT, Fig. 7B3), glycine decarboxylase complex and serine hyrdroxymethyltransferase (GDC and SHMT, Fig. 7B4)) (Peterson, 1982; Havir, 1986; Lu et al., 2014). β-oxidation and regeneration of the glyoxylate cycle substrate oxaloacetate from succinate could also elevate cellular redox potential leading to NADH inhibition of GDC in photorespiration (Fig. 7A4-5, 7-9; 7B1-5) (Igamberdiev and Lea, 2002; Ma et al., 2016; Igamberdiev and Bykova, 2018; Igamberdiev, 2020). Further, if GDC were inhibited in the mitochondria, this could result in accumulation of photorespiratory metabolites that modulate Calvin cycle and photorespiration (Fig. 7A9; 7B4-5) (Peterson, 1982; Igamberdiev et al., 2004; Timm et al., 2012; Bykova et al., 2014; Negi et al., 2018). Some support for impaired photorespiration is available from previous transcriptomic and metabolomic analysis of HO compared to WT that suggested increased expression of RuBisCO, GDC, SHMT, glutamine synthetase (GS), and hydroxy pyruvate reductase (HPR) and altered levels of glycine, glycerate, 2-oxoglutarate (2-OG) (Vanhercke et al., 2017; Mitchell et al., 2020). Altered metabolite and transcript levels are consistent with photorespiratory phenotypes previously described (Timm and Bauwe, 2013). Under normal circumstances the glyoxylate cycle and photorespiration are thought to be developmentally separated to germinating seeds and leaves, respectively; however, in HO leaves the combined presence could explain the ∼50% reduction in CO_2_ fixation and adverse growth phenotypes.

### Hypothesis 2: TAG turnover feeds starch synthesis and distorts sink strength leading to reduced CO_2_ assimilation in the HO line

The cycling of ^14^C between TAG and starch in HO (Fig. 6C-D) suggests carbon from TAG production and turnover is capable of partially offsetting photosynthetically derived starch synthesis (Fig. 7A4-6) if enzymatic steps in gluconeogenesis were active. Specifically, the initial limited partitioning of fixed ^14^CO_2_ to starch in HO (Fig. 6, 0 hrs) may be a consequence of TAG catabolism during this time (Fig. 7A5-6) which would provide an unlabeled source of carbon for starch. Use of TAG this way could reduce photosynthetic demand for starch, resulting in decreased photosynthetic carbon uptake as has been shown for starchless mutants (Huber and Hanson, 1992; Edwards et al., 1999). A starchless *Arabidopsis thaliana* mutant was improved (i.e., carbon uptake and growth) by directing carbon flux from sugars to lipids and suggests lipids synthesis can be leveraged to increase carbon sink strength (Fan et al., 2019). Furthermore, when lipid storage capacity was enhanced in ryegrass, the authors also observed increased photosynthesis and total carbon content (Beechey-Gradwell et al., 2020). Both studies are similar to the LEC2 line where enhanced fatty acid synthesis and TAG stability (Fig. 5) correlated with enhanced CO_2_ fixation (Fig. 3A). The shift in carbon flux to a TAG-starch futile cycle may reflect recently predicted targets of WRI1 in the upper glycolysis and pentose phosphate pathways (Kuczynski et al., 2022) whereby WRI1, also involved in the regulation of cell wall biosynthesis (Haigler et al., 2009; Qu et al., 2012; Qaisar et al., 2017) may account for the unexpected increase of ^14^C in the HO cell wall (Fig. 3D). These reports support the assertion that altered carbon partitioning and TAG catabolism negatively modulates carbon fixation in photosynthetically active leaves engineered to accumulate lipids. The reassimilation of carbon from TAG turnover into starch biosynthesis reduces the sink strength of starch for photosynthetic carbon to further limit photosynthetic carbon capture.

### Hypothesis 3: *AtLEC2* stabilizes TAG accumulation preventing the HO TAG-starch futile cycle and increasing CO_2_ assimilation and plant growth

Compared to the first-generation HO tobacco line, the LEC2 accumulated twice as much TAG while mostly alleviating the growth defects of the HO (Vanhercke et al., 2014; Vanhercke et al., 2017). Previously, the metabolic causes for these differential phenotypes were unclear. Here we demonstrate that the increase in leaf oil accumulation in LEC2 did not come from further diversion of membrane lipids to TAG (Fig. 1), and the enhanced growth was likely associated in part with recovered ^14^CO_2_ fixation (Fig. 3A). Interestingly partitioning of “new” [^14^C]carbon to the lipid and TAG fractions increased for LEC2 relative to HO (Fig. 3F; Fig. 5) and this increase was attributed to a combination of new carbon partitioning to lipid synthesis and improved TAG stability by reducing TAG degradation in the LEC2 line (Fig. 1, Fig. 5). Recent evidence suggests that co-expressing *LEC2* and *WRI1* promotes TAG synthesis and stability beyond that of *WRI1* alone through enhanced expression of fatty acid synthesis and oil body packing (OLE1/OLE2/OLE3/OLE4/OLE5) (Baud et al., 2007; Kim et al., 2015). We suggest the metabolic stability of TAG in the LEC2 line (Fig. 5) limited the detrimental effects of the TAG-starch futile cycle described in Hypothesis 1 and 2, and thus contributed to the enhanced growth and carbon capture of LEC2 over HO.

## Conclusion

The ability to greatly increase the production of plant oils per area of land through the engineering of novel vegetative oil crops holds great potential, but both the amount of oil produced and the effects on plant growth have varied widely in various crop plants (Vanhercke et al., 2019). Utilizing two previously produced oil accumulating tobacco lines we describe the stability of engineered leaf oil accumulation is key to limited adverse metabolic responses in central carbon metabolism (Fig. 7) that otherwise reduce CO_2_ assimilation and growth. An inefficient TAG-starch futile cycle in HO that is alleviated in the LEC2 line likely accounts for much of the growth and metabolic differences between the lines. Together with a recent report that tobacco accumulates non-transient starch in leaves which can provide the excess carbon for lipid synthesis (Chu et al., 2022), the results suggest the capacity of tobacco to capture more CO_2_ than needed for vegetative growth and store it as a metabolically stable product (non-transient starch, or TAG in LEC2) may account for the success in tobacco leaf oil engineering relative to other species. Interestingly, while the LEC2 is similar to WT in size, ^14^CO_2_ assimilation was 83.3% of WT suggesting further engineering of the LEC2 line may be able to enhance metabolic performance and further increase leaf oil accumulation.

## METHODS

### Plant growth

Tobacco seeds were germinated in a growth chamber at 16 h light/8 h dark, 23°C, and 100-300 µmol m^-2^ s^-1^ light at pot level (3.5 inch pots). At 30 days after sowing (DAS), plants were transferred to 2.8-liter pots in a greenhouse with supplemental lighting and heating set to 16 h light/8 h dark, 28*°C*/20*°C, and* 200-400 µmol photons m^−2^ s^−1^. Watered daily, as needed, and fertilized twice per week with 20/10/20 NPK (1 part fertilizer: 15 parts water) with an additional 200 ppm micronutrient solution of Scotts Miracle-Gro.

### Leaf lipid mass measurements

Three plants utilized per genotype for total fatty acid analysis, 7 mm diameter leaf disks were collected from 1-3 leaves at 30-DAS, 16 mm disks were collected from 4-6 leaves at 44-DAS, and from 5-7 leaves at 64-DAS. Disks were submerged in 1.5 mL 2.5% sulfuric acid in methanol (v/v) in glass tubes containing 17:0 TAG standard and incubated at 80°C for 1 hr to produce fatty acid methyl esters (FAME). 0.5 mL hexanes and 3 mL 0.9% KCl (w/v) was added to induce phase separation. FAME in the upper hexanes phase concentrated under N_2_ and re-suspended in 0.2 mL hexanes for quantification via gas chromatography with flame ionization detection (GC-FID) on Restek FATwax column (30 m, 0.25 inner diameter, 0.25 mm film thickness). 5 µL injection, 1/40 split ratio, 0.9 ml/min helium flow rate. Heating conditions: 170 °C ramped 10 °C per minute to 230 °C, hold 5 minutes. FAME data was normalized to a leaf disk area. For lipid class analysis, lipid extraction and thin layer chromatography (TLC) were done as previously (Zhou et al., 2020). Lipid classes identified by TLC migration with standards after staining with 0.05% primulin in acetone/water 80:20 (v/v) and visualized under UV light, then removed from TLC for conversion to FAME and analyzed by GC-FID.

### ^14^CO_2_ Pulse chase

35 DAS plants were transferred to the lab for 10 days to acclimate to environmental conditions of ∼350 µmol photons m^−2^ s^−1^ of fluorescent light at 22°C-30°C and ∼20-30% humidity. The 45-day-old tobacco plants (2 WT, 3 HO, 3 LEC2) were placed together in a 65-L desiccator under the lights containing two small fans, and a vial of 1 mCi [^14^C]bicarbonate (American Radiolabeled Chemicals, Inc). The 2 hr pulse (evolution of ^14^CO_2_) was triggered by delivering 2 mL of 5 M sulfuric acid to the [^14^C]bicarbonate via tubing and syringe at 5 hours into the day. At the end of the pulse the chamber was purged inside a fume hood through a 1 M KOH base trap to capture any remaining ^14^CO_2_. Once removed the plants were placed under lights and the 0 hr leaf disks were collected from the first three sequential leaves above the cotyledons near the tip (12-WT, 18-HO, 18-LEC2 replicates). T For subsequent time points, two disks were collected from each side of the midrib moving toward the base 4-WT, 6-HO, 6-LEC2 replicates (Fig. 2A). All leaf disks were immediately frozen in liquid N_2_ and stored at −80°C.

### Extraction and Analysis of ^14^C Fractions

All solvents and tools were stored at −20°C or on ice until use. Leaf disks were homogenized by bead beater in 2 mL tubes with five 2.4 mm ceramic beads for 3-4 intervals of 30 seconds on medium speed. After each interval, samples were frozen in liquid nitrogen for 20 seconds. The extraction of lipids, organic acids, amino acids, sugars, protein, starch, and cell wall was as in (Allen and Young, 2013) except for modifications of chloroform/methanol/1 M formic acid (20/10/1; v/v/v) for lipid extraction solvent, and a 3 hr Solusol (National Diagnostics) digestion to dissolve the cell wall fraction. Radioactivity in aliquots of each fraction was quantified with a Packard Tri-Carb 2200 scintillation counter and data analyzed with Microsoft Excel and R. An aliquot of the labeled lipid fraction was also used to measure total mass by GC-FID of FAME and individual lipid classes by TLC as above.

## ACKNOWLEDMENTS

We thank Xue-Rong Zhou of CSIRO, Australia for providing the transgenic tobacco lines.

## AUTHOR CONTRIBUTIONS

P.D.B. and D.K.A. conceived the original research plans and acquired the funding; B.S.J. performed the experiments; all authors analyzed the data and contributed to writing the article.

